# An *in vitro* Chondro-osteo-vascular Triphasic Model of the Osteochondral Complex

**DOI:** 10.1101/2020.08.27.270660

**Authors:** Alessandro Pirosa, Riccardo Gottardi, Peter G. Alexander, Dario Puppi, Federica Chiellini, Rocky S. Tuan

## Abstract

The generation of engineered models of the osteochondral complex to study its pathologies and develop possible treatments is hindered by the distinctly different properties of articular cartilage and subchondral bone, with the latter characterized by vascularization. *In vitro* models of the osteochondral complex have been mainly engineered as biphasic constructs containing just cartilage and bone cells, a condition very dissimilar from the *in vivo* environment. The different cellular components of the osteochondral complex are governed by interacting biochemical signaling; hence, to study the crosstalk among chondrocytes, osteoblasts, and endothelial cells, we have developed a novel triphasic model of the osteochondral tissue interface. Wet-spun poly(ε-caprolactone) (PCL) and PCL/hydroxyapatite (HA) scaffolds in combination with a methacrylated gelatin (gelMA) hydrogel were used as the polymeric backbone of the constructs. The scaffold components were engineered with human bone marrow derived mesenchymal stem cells (hMSCs) and human umbilical vein endothelial cells (HUVECs), and differentiated using a dual chamber microphysiological system (MPS) bioreactor that allows the simultaneous, separate flow of media of different compositions for induced differentiation of each compartment towards a cartilaginous or osseous lineage. Within the engineered Microphysiological Vascularized Osteochondral System (microVOCS), hMSCs showed spatially distinct chondrogenic and osteogenic markers in terms of histology and gene expression. HUVECs formed a stable capillary-like network in the engineered bone compartment and enhanced both chondrogenic and osteogenic differentiation of hMSCs, resulting in the generation of an *in vitro* system that mimics a vascularized osteochondral interface tissue.

## 1 Introduction

The osteochondral (OC) interphase tissue represents a major target of degenerative joint diseases, such as osteoarthritis (OA), a primary cause of physical disability in millions of people worldwide, e.g., up to 60 million people in the United States [1]. OA affects the whole joint but is primarily characterized by the progressive degeneration of articular cartilage and subchondral bone dysfunction. There are currently no available treatments except palliative ones, ultimately requiring total joint replacement when pain and loss of function become unmanageable. The crosstalk between articular cartilage and subchondral bone is generally considered to play a major role in disease progression, but the molecular mechanisms underlying this crosstalk and OA pathogenesis are incompletely understood.

Although the progression of OA results from the coordinated dysfunction between all tissues of the synovial joint, the etiology of osteoarthritis is well-defined only in the case of post-traumatic osteoarthritis [2], a clear case of mechanical joint surface injury leading to joint degeneration. However, bone is also involved in post-traumatic OA. Bone bruises seen on magnetic resonance imaging following anterior cruciate ligament injuries are very common [3] and a significant relationship between area of initial bone bruise at the time of injury and progressive posttraumatic chondral disease has been reported [4–6]. The bruising is caused by a disruption in the blood flow of the cancellous bone, that is resolved in an as-yet-unidentified manner, presumably involving interactions between osteoblasts, endothelial cells, and mesenchymal stem cells, among others [7] and may result in altered mechanics and osteochondral crosstalk that ultimately contribute to OA.

Co-culture systems have been used to study the interactions between cell populations, such as chondrocytes and osteoblasts, but they lack the fundamental three-dimensional (3D) and mechanical cues typical of the OC complex, especially since this interface is subjected to load-bearing conditions [8]. Tissue engineering approaches have therefore been developed to produce biomimetic constructs that mimic the anatomical and physiological hierarchical structure of the OC complex [9]. The most studied constructs for OC engineering are represented by bilayered or gradient composite structures with different local mechanical, structural and chemical properties [10,11]. These models have allowed researchers to elucidate a number of biochemical interactions between bone and cartilage cells in the pathogenesis of joint diseases such as OA. Findings include the production by subchondral bone of cytokines and growth factors promoting loss of cartilage proteoglycans [12], as well as the increased synthesis by chondrocytes of matrix degrading molecules such as metalloproteinases and proinflammatory cytokines [13]. However, these studies did not take into account the contribution of other cell types, such as endothelial cells. Vascular infiltration from the subchondral bone to the articular cartilage has been documented in the progression of OA [14–16].

The role of vasculature in bone and osteochondral development, growth, and repair is well documented. Vascular invasion is required for endochondral ossification and the ossification of cartilage templates in fetal skeletal development [17], regulation of growth plate maturation and long bone growth, as well as cortical bone fracture repair where the cartilaginous callous serves as template for the new bone [18,19]. Furthermore, vasculature has been found to promote expression of osteogenic genes in hypertrophic chondrocytes, thus promoting initiating chondrocyte-to-osteoblast transformation [20], and vasculature introduces osteoblasts to complete bone formation. Even during intramembranous ossification, where cartilaginous structures are not maintained, vascularization is still required for delivery and transport of nutrients, metabolites, and cells [7]. Importantly, the vasculature does not serve only as conduits, but also as sources of cytokines that enhance bone repair [21].

Altered crosstalk between osteoblasts and endothelial cells has also been shown to be related to the pathogenesis of several bone diseases, such as osteomyelitis, avascular necrosis and osteoporosis [22]; however little is known about the interaction of bone with cartilage across the subchondral plate in the development of these pathologies and osteoarthritis. All these phenomena are regulated by precise cellular and molecular interactions, the elucidation of which would certainly inform the development of OA treatments and fracture healing augmentation.

The generation of an experimental model of the OC complex that comprises an endothelial component is thus crucial to better understanding the development and pathologies of the OC complex. The majority of *in vivo* vascularization approaches use angiogenic factors combined with 3D scaffolds [23,24] and pre-vascularization strategies [25], whereas the use of co-culture systems combined with 3D scaffolds has been mostly used for engineering *in vitro* constructs [26,27]. One example of using a co-culture system to introduce osteogenic differentiation and vasculature development *in vitro* was made by Tsigkou *et al.*, which combined human bone marrow-derived mesenchymal stem cells (hMSCs) and human umbilical vein endothelial cells (HUVECs) seeded in a polymeric scaffold and a hydrogel, respectively. They observed the formation of HUVECs’ capillary-like structures 4-7 days after implantation into a mice model, while the hMSCs were necessary for the development of a stable vasculature [26].

Here, we report the development of a novel *in vitro* vascularized OC construct to recreate the triphasic nature of the OC complex. We utilized a microphysiological system (MPS) bioreactor we previously designed that maintains two separate compartments for the chondral and osseous differentiation within a contiguous construct [28,29]. The cartilage construct was engineered by incorporating hMSCs in a photocrosslinkable methacrylated gelatin (gelMA) hydrogel [30], while the osseous construct was generated by seeding hMSCs on a poly(ε-caprolactone) (PCL) and PCL/hydroxyapatite (HA) scaffold, produced by an additive manufacturing technique previously developed by Puppi *et al.* [31]. The wet-spun scaffold aims to mimic the mechanically stiffer and more porous osseous support, while the gelMA hydrogel mimics the characteristics of the cartilage extracellular matrix (ECM). A capillary-like network is introduced in the PCL-PCL/HA scaffolds by using HUVECs, exploiting the wide porosity of the wet-spun scaffolds. This engineered microphysiological vascularized osteochondral system (microVOCS) provides a biomimetic system that recapitulates the morphophysiology of the OC complex, and can be employed as a platform for the study of OC diseases and the testing of pharmacological treatments.

## 2 Materials and Methods

### 2.1 Materials

Poly(ε-caprolactone) (PCL, CAPA 6500, molecular weight = 50,000) was supplied by Perstorp Caprolactones Ltd (Warrington, UK). HA nanoparticles (size < 200 nm) were purchased from Sigma-Aldrich (Milan, Italy). Gelatin was purchased from Sigma-Aldrich (St. Louis, MO).

### 2.2 Bioreactor Design and Fabrication

The 3D structure of the bioreactor was modeled using Solidworks (Waltham, MA) and all components were fabricated using a 3D Systems Viper (Rock Hill, SC) and WaterShed XC 11122 resin (DSM Somos®, Heerlen, Netherlands) [32].

### 2.3 Preparation of PCL and PCL/HA scaffolds

PCL and PCL/HA scaffolds were produced by computer-aided wet-spinning (CAWS) as described elsewhere [31,33,34]. Briefly a PCL solution 20 % w/v in acetone (with HA nanoparticles added at 1:4 w/w ratio to the polymer for the PCL/HA scaffolds) was placed into a glass syringe fitted with a stainless steel blunt needle (internal diameter = 0.41 mm) and extruded in an ethanol coagulation bath using a computer-controlled rapid prototyping machine (MDX-40A, Roland DG Mid Europe Srl, Italy). An initial distance between the needle tip and the bottom of the beaker (Z_0_) of 3 mm and a flow rate of 1 mL/h were set in all the performed experiments. The 3D geometrical scaffold parameters, including the distance between fiber axis (d_xy_), layer thickness, scaffold external geometry and sizes, were designed using an algorithm developed in MATLAB software (The Mathworks, Inc.). The combination of the X–Z axis needle motion and the Y axis platform motion was used to fabricate scaffolds layer-by-layer. A biopsy puncher (Kai Industries, Japan) with an internal cutting diameter of 4 mm was used to obtain four cylindrical samples from a square-based template scaffold with side of about 15 mm and height of about 5 mm. The morphological properties of the scaffolds were evaluated using an Olympus SZX16 stereomicroscope (T2 TOKYO, Japan) and dimensional analysis was performed using ImageJ software (NIH).

### 2.4 Preparation of gelatin methacrylated (gelMA)

GelMA was produced by the reaction between gelatin and methacrylic anhydride (MA) in water according to a previously described procedure [35]. Briefly, type A gelatin was dissolved in deionized water at 37 °C and mixed with methacrylic anhydride for 12 h, to allow the reaction between amine side groups of gelatin and carboxyl groups of MA. The reaction mixture was dialyzed for 4 days against distilled water at room temperature and the reaction product was lyophilized and stored in a desiccator. The photoinitiator lithium phenyl-2,4,6-trimethylbenzoylphosphinate (LAP) was synthesized as previously described by Fairbanks *et al*. [36]. The liquid gelMA was photocrosslinked by UV light for 2 minutes.

### 2.5 Cell culture

hMSCs were extracted from femoral heads of patients undergoing total joint arthroplasty with IRB approval (University of Pittsburgh), according to the procedure reported by Caterson *et al*. [37]. hMSCs were expanded as monolayer culture in Dulbecco’s Modified Eagle Medium (DMEM) supplemented with 10 % fetal bovine serum and 2 % penicillin/streptomycin/fungizone (growth medium, GM) at 37 °C and 5 % CO_2_. GM was changed every 2 ‒ 3 days until the cells reached ~80 – 90 % confluency. Experiments were conducted on cells extracted from nine different patients divided into three pooled groups of three (n = 3). Lentiviral-GFP transfected HUVECs (from now on referred to as HUVECs) were purchased from Angio-Proteomie (Boston, MA) and grown as monolayer in Endothelial Growth Basal Medium 2 (EBM-2) supplemented with the EGM-2 Bullet kit, both purchased from Lonza (Basel, Switzerland).

### 2.6 Sterilization and conditioning of the scaffolds

Scaffolds were sterilized in a 70 % ethanol aqueous solution for three hours, dried under sterile biosafety cabinet for 1 h, extensively washed with phosphate buffered saline (PBS) to remove residual ethanol and then rinsed with complete GM. Scaffolds were then incubated at 37 °C in GM at 5 % CO_2_ until cell seeding.

### 2.7 Fabrication of osteochondral constructs

<u>Preparation of osseous constructs</u>. PCL-PCL/HA scaffolds were seeded with 100 μl of hMSCs suspension at a concentration of 8×10^5^ cells/mL. A 25 μl aliquot of cell suspension was gently poured on four different sides of the scaffolds every 30 minutes, to achieve uniform distribution of cells within the samples. After cell seeding, scaffolds were submerged in GM and cultured for 10 days before insertion into the bioreactor. <u>Preparation of osteochondral constructs</u>. After 10 days of proliferation, PCL-PCL/HA scaffolds were placed in the bottom chamber of the bioreactor insert, which was then inserted into the bioreactor well. For the chondral component, hMSCs were resuspended in the 10 % gelMA/0.15 % LAP (w/v) PBS solution (pH 7.4) at a final density of 1×10^7^ cell/mL, and then poured onto the scaffold to fill the upper part of the insert and then irradiated with UV light for 2 min to obtain a solid gelMA hydrogel. Chondrogenic medium (CM) was supplied through the upper conduit, while osteogenic medium (OM) through the bottom conduit. The following compositions were used for the differentiation media: (1) OM = GM supplemented with 0.1 mM L-ascorbic acid-2 phosphate, 10 mM β-glycerophosphate and 10 nM 1,25-dihydroxyvitamin D_3_; (2) CM = GM without FBS supplemented with 10 ng/mL TGF-β3, 1 % insulin-transferrin-selenium, 50 μM L-ascorbic acid-2 phosphate, 10 nM dexamethasone and 23 μM proline. The perfusion rate was 1 mL/day, and the medium reservoirs were replenished every 9 days. After 4 weeks of differentiation, the engineered OC constructs were collected for validation using histological/immunohistochemical (IHC) analysis and quantitative reverse transcription real-time PCR (qRT-PCR).

### 2.8 Fabrication of vascularized osteochondral constructs

<u>Preparation of vascularized osseous constructs</u>. PCL and PCL/HA scaffolds were seeded and cultured with hMSCs as described in paragraph §2.7. After 2 weeks of osteogenic differentiation *in situ*, a mixture of hMSCs:HUVECs (1:4 ratio) was resuspended in 10 % gelMA/0.15 % LAP (w/v) PBS solution diluted 1:1 with EGM-2 (final gelMA concentration 5 % w/v) at a final density of 1×10^6^ cell/mL. The cell suspension was poured into the PCL and PCL/HA scaffolds and allowed to penetrate and fill the scaffold pores. Subsequently, the suspension was gelated by irradiation for 2 min with UV light. Constructs were incubated with OM:EGM-2 1:1 (v/v) for two more weeks to obtain a vascularized osseous construct. <u>Preparation of vascularized osteochondral constructs</u>. After 2 weeks of osteogenic differentiation in the lower chamber of the bioreactor, a suspension of hMSCs:HUVECs (1:4 ratio) was prepared as described previously and poured onto the PCL and PCL/HA scaffolds. Subsequently, the suspension was gelated by irradiation for 2 min with UV light to obtain a triphasic construct. Constructs were cultured with OM:EGM-2 1:1 (v/v) in the lower flow stream and CM in the upper flow stream for two more weeks.

### 2.9 Live/Dead assay

Viability of HUVECs and hMSCs was assessed using a Live/Dead assay, staining cells with Calcein Blue AM (ex/em ~ 360/445 nm) (eBioscience, USA) and ethidium homodimer (EthD-1, ex/em ~ 495/635 nm) (Molecular Probes, USA), for the detection of live and dead cells, respectively. Cell-seeded constructs were submerged in the Calcein Blue AM/EthD-1 (4 μM/8 μM) solution for 45 min at room temperature in the dark, and then observed using an Olympus SZX16 fluorescence stereomicroscope (T2 TOKYO, Japan).

### 2.10 Histology and immunohistochemistry (IHC)

Samples were fixed in 4 % paraformaldehyde in PBS solution, paraffin embedded following standard procedures and sectioned at 20 μm thickness. Sections were re-hydrated and stained with alcian blue and alizarin red to detect glycosaminoglycans (GAGs) and calcium deposits, respectively. Hematoxylin/Eosin (H&E) staining was used to observe cell morphology. For IHC, heat-mediated antigen retrieval was performed with sodium citrate buffer (eBioscience, USA) at pH 6.0 for 20 min at 90 °C. Endogenous peroxidase activity was quenched by incubating in 3% H_2_O_2_ solution in methanol (10 min treatment), while nonspecific binding was blocked with 1% horse serum (Vector Labs) in PBS at room temperature for 45 min. After antigen retrieval and blocking, samples were incubated overnight at 4 °C with primary anti-human CD31 antibodies (Abcam, cat. n° ab28364) or anti-human pAKT (Cell Signaling, cat. n° 4060) antibodies at a dilution of 1:100 and 1:50 respectively, following by an incubation of 30 min with biotinylated secondary antibodies (Vector Labs). Staining was achieved using horseradish peroxidase (HRP)-conjugated streptavidin and Vector® NovaRED™ peroxidase substrate, with counterstaining performed with hematoxylin (Vector Labs). Following staining, sections processed for histology and IHC samples were dehydrated, mounted and coverslipped, and images were acquired with a Nikon Eclipse E800 microscope (Nikon Instruments Inc., Melville, NY).

### 2.11 Quantitative reverse transcription real-time PCR

The osseous and chondral compartments were detached at the experimental end-point and processed separately for analysis. Total RNA was extracted using Trizol (Invitrogen, USA) and purified with the RNeasy Plus mini kit (Qiagen, Germany) following the manufacturers’ protocols. Reverse transcription was performed using the SuperScript IV kit (Invitrogen, USA) utilizing random hexamers for the preparation of cDNA. Real-time PCR was performed using the StepOnePlus thermocycler (Applied Biosystems, CA) and SYBR Green Reaction Mix (Applied Biosystems, CA). Human osteopontin (OPN), osteocalcin (OSC) and bone sialoprotein 2 (BSP2) were analyzed to assess osteogenic differentiation; aggrecan (ACAN), SRY (sex determining region Y)-box 9 (SOX9) and collagen type II (COL2) were analyzed to assess chondrogenic differentiation; and endothelin-1 (ET-1) was analyzed for HUVECs activity. hMSCs or hMSCs/HUVECs seeded in the scaffolds and cultured in GM after 10 days of proliferation were used as “day 0” for osteogenesis; cells resuspended in the gelMA solution prior to differentiation were used as “day 0” for chondrogenesis. Transcript level of 18S rRNA was used as endogenous control, and gene expression fold of change was calculated by the comparative cycle threshold (CT) method, using expression levels at day 0 as reference for the 2^−ΔΔCT^ calculation.

**Table 1.**
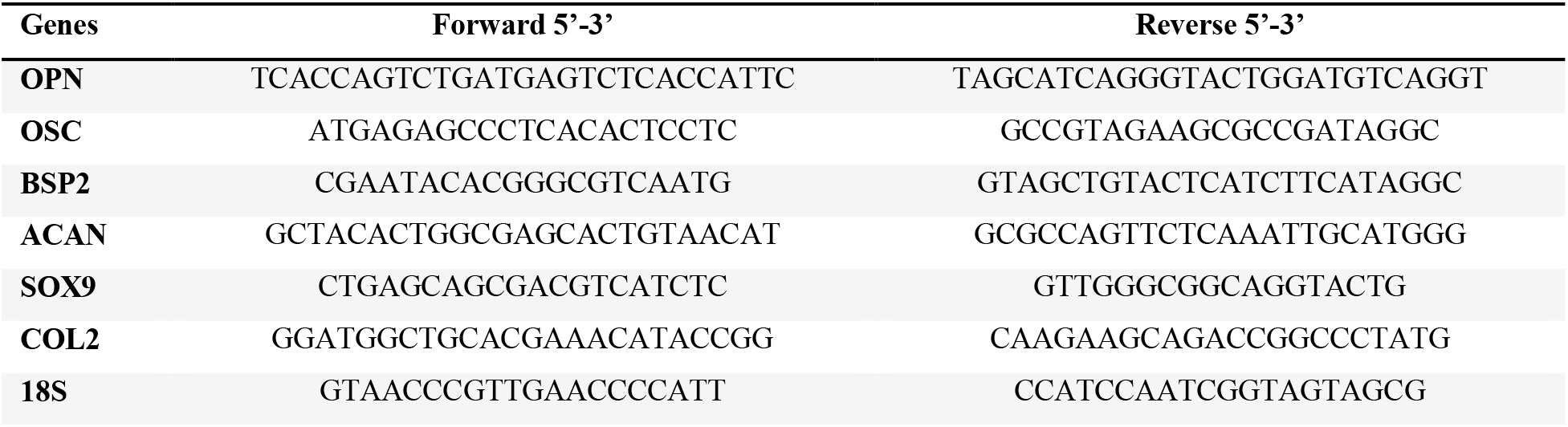
PCR primer sequences for qRT-PCR analysis of gene expression

### 2.12 Statistical analysis

Each sample was assayed in triplicate and quantitative data were reported as mean ± SD. Statistical analyses were performed using GraphPad Prism 7 (GraphPad Software Inc., San Diego, CA). Student’s *t*-test and one-way ANOVA/post hoc Tukey test were used to compare two or more independent groups, respectively. Statistical tests were two-tailed and significance was set at p-value < 0.05.

## 3 Results

### 3.1 HUVECs enhance osteogenesis in the developed vascularized bone constructs

PCL and PCL/HA scaffolds for the development of the bone compartment were produced by computer-aided wet-spinning (CAWS), which involved the extrusion of a polymeric solution through a X-Y-Z translating needle that was immersed into a coagulation bath [31] (Figure 1a).

**Figure 1.**
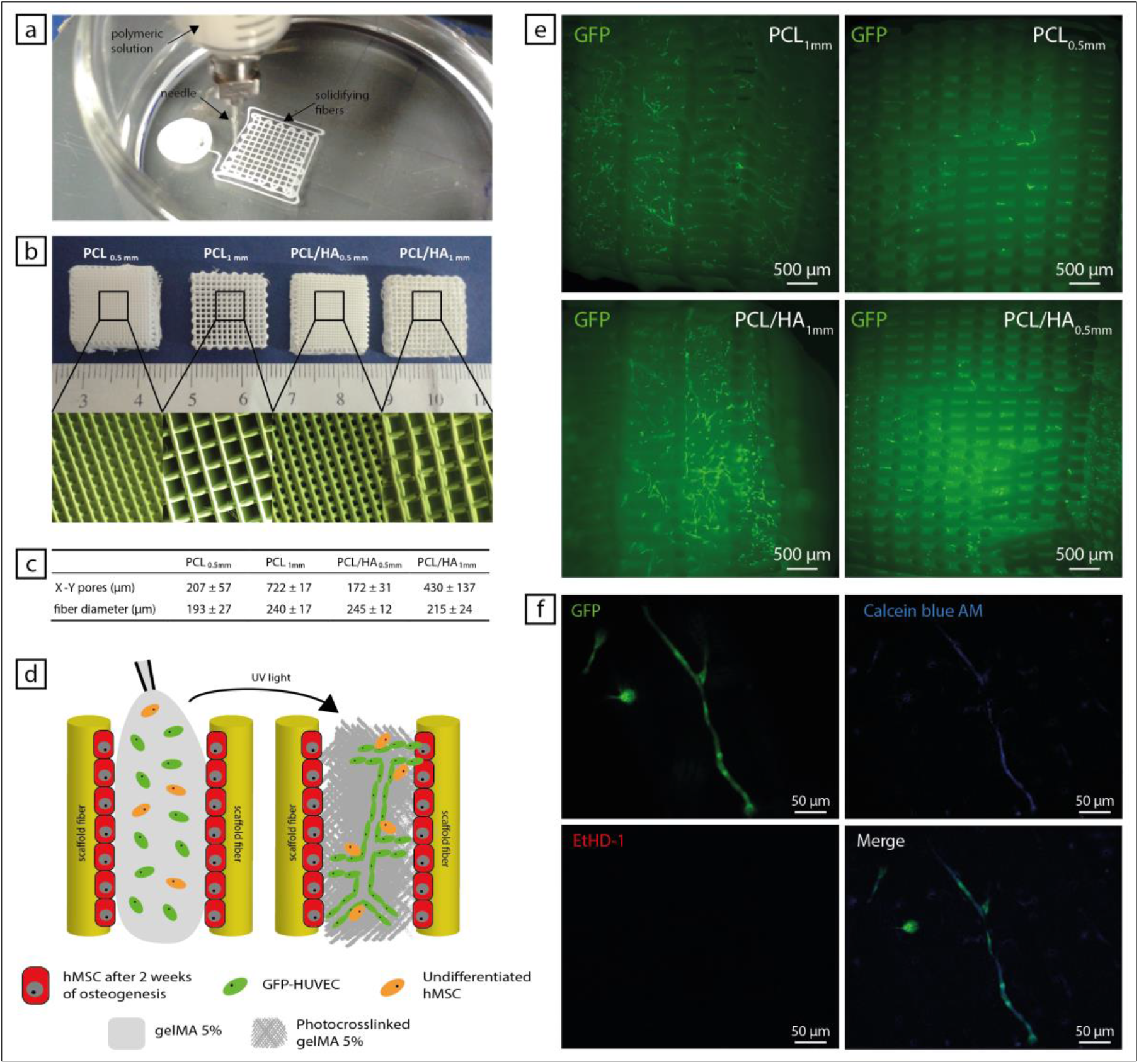
Vascular network formation in 3D printed osseous scaffolds. (a) Production by CAWS of the scaffolds; the polymeric solution inside the syringe was injected in the ethanol coagulation bath with a 0-90 ° lay-down pattern. (b) Macroscopic images and stereomicroscope magnifications of the scaffolds produced with different X-Y inter-fiber axes distances and compositions; scale bar = 1 mm. (c) Morphological parameters of the scaffolds obtained by ImageJ software. (d) Schematic of the preparation of the vascularized bone constructs. (e) Representative HUVEC-derived capillary-like structures inside PCL and PCL/HA scaffolds combined with gelMA. (f) Representative Live/Dead assay of HUVECs/hMSCs co-culture on PCL1mm-gelMA construct: GFP = HUVECs; Calcein Blue AM = live cells (blue); EtHD-1 = dead cells (red).

A layer composed of parallel fibers was fabricated by depositing the solidifying solution filament with a predefined pattern, and 3D architectures were built layer-by-layer by fabricating layers with alternating fiber orientation (0 – 90° lay-down pattern) one on top of the other (Figure 1a). Two different scaffold architectures were developed (Figure 1b): one was obtained with an X-Y inter-fiber needle translation distance (d_xy_) of 0.5 mm (PCL_0.5mm_ and PCL/HA_0.5mm_); the other was obtained with a d_xy_ of 1 mm (PCL_1mm_ and PCL/HA_1mm_). All scaffolds possessed a well-defined fibrous structure, composed of aligned fibers with a 0 ‒ 90 ° lay-down pattern. Morphological parameters of the produced scaffolds are reported in Figure 1c. HUVECs were used to enhance the osteogenic potential of hMSCs and represented the endothelial component of our model. Scaffolds were seeded with hMSCs and cultured under osteogenic differentiation for 2 weeks. Thereafter, a mixture of HUVECs/hMSCs at a 4:1 ratio resuspended in 5 % gelMA (10 % gelMA:EGM-2 1:1 v/v) and poured into the scaffolds and allowed to infiltrate via the interconnected pores (Figure 1d). The constructs were irradiated with UV-light for 2 minutes to photocrosslink the gelMA and obtain a hydrogel network encapsulating HUVEC/hMSCs. HUVECs formed interconnected capillary-like structures within the 5 % gelMA in the PCL_1mm_ and PCL/HA_1mm_ scaffolds throughout all the pores, as revealed by GFP-positive and CD31-positive signals (Figure 1e and Figure 2a). PCL_0.5mm_ and PCL/HA_0.5mm_ scaffolds did not show endothelial tubular formation, probably due to the lack of gelMA penetration within the smaller sized pores. For this reason, PCL_0.5mm_ and PCL/HA_0.5mm_ scaffolds were not used for subsequent experiments. Live/Dead analysis of HUVEC/hMSC co-culture on PCL_1mm_-gelMA construct revealed the presence of viable hMSCs (blue only stained cells in Figure 1f) supporting HUVEC-derived capillary-like structures (Figure 1f, green and blue stained cells). There were no dead cells, as indicated by the absence of red stained cells, suggesting adequate nutrient transport within the mature scaffolds, even in the presence of gelMA filling the interior pores. Histological analysis revealed more calcium deposits in the constructs containing the HUVECs (Figure 2a), confirming their positive effect on the osteogenic potential of hMSCs, as reported in other studies [26,38].

**Figure 2.**
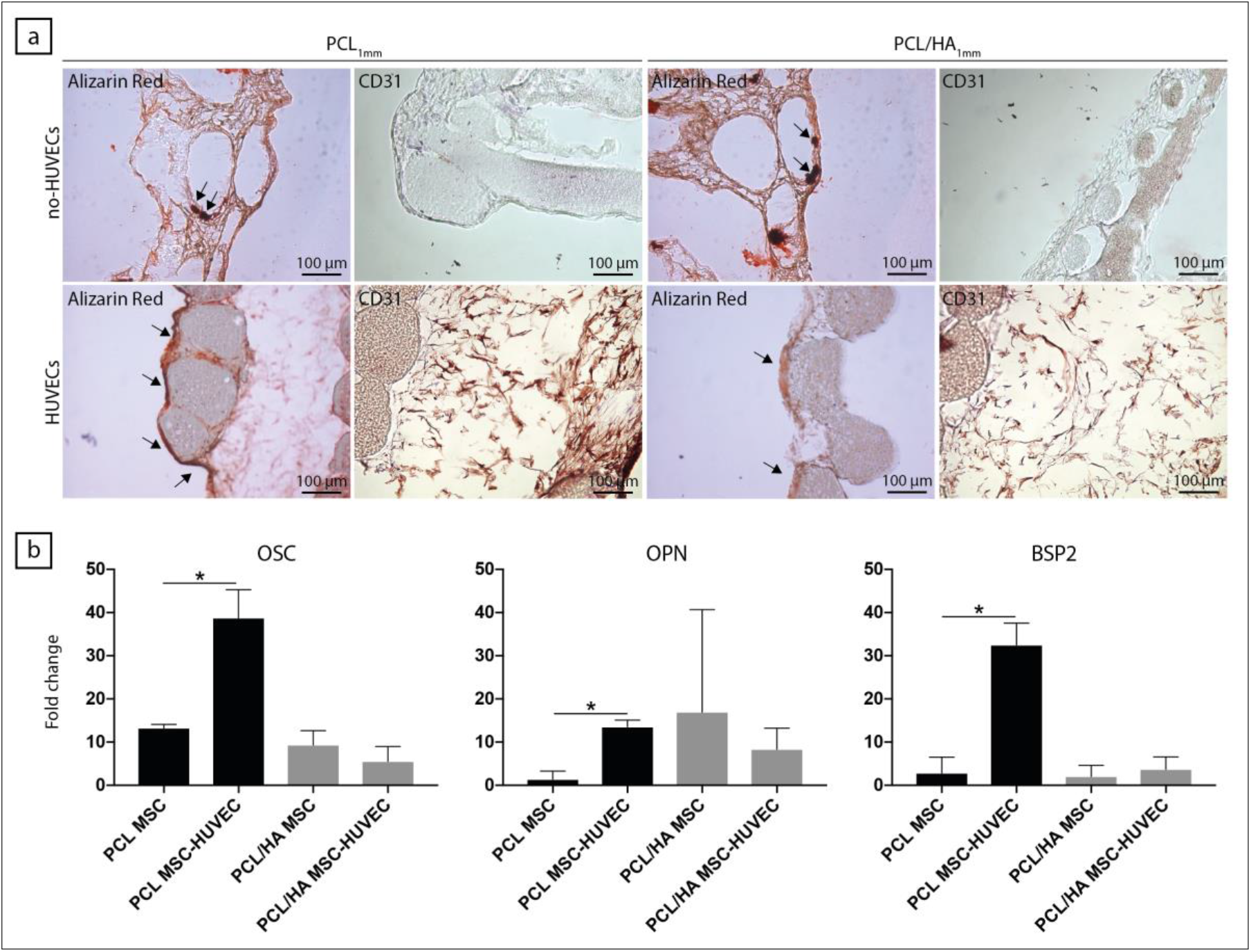
The effect of HUVECs on hMSCs osteogenesis in osseous scaffolds with and without nanohydroxyapatite. (a) Representative histological and CD31-IHC images of PCL-PCL/HA scaffolds seeded with hMSCs and HUVECs/hMSCs. Arrows: calcium deposits detected by intense alizarin red staining. CD31 positive red staining indicate the presence of HUVECs. (b) Real time qRT-PCR of osteogenic genes showing increased osteogenic differentiation of hMSCs in the presence of HUVECs. Fold change = 1 represents baseline of undifferentiated cells; * p < 0.05.

Gene expression profiles of hMSCs in mature PCL and PCL/HA scaffolds in the presence or absence of HUVECs were evaluated after 2 weeks of osteogenic differentiation to quantify the effects of vascularization on hMSCs osteogenic commitment. As shown in Figure 2b, in the PCL scaffolds, expression of osteogenic markers (OSC, OPN and BSP2) was up-regulated in the presence of HUVECs in comparison to hMSCs alone, confirming the enhancement of hMSC osteogenic potential in the presence of an endothelial microvascular network [38,39]. With hydroxyapatite (HA) present in the scaffold, the HUVECs-mediated enhancement of osteogenic differentiation was lower, with no positive effect on OPN and OSC expression.

### 3.2 MPS bioreactor for the development of in vitro biphasic osteochondral constructs

The external geometry and dimensions of the PCL and PCL/HA scaffolds were tailored for culture within a dual chamber MPS bioreactor we previously developed and characterized for osteochondral differentiation and disease modeling [40,41]. After extracting cylindrical cores from the template scaffold, hMSCs were allowed to proliferate on the PCL-PCL/HA scaffolds in the presence of growth medium for 10 days to assure optimal colonization. The cell-seeded scaffolds were then placed in the bottom chamber of the insert, while gelMA containing undifferentiated hMSCs at a density of 1×10^7^ cells/mL was poured to fill the top chamber and photocrosslinked by irradiation with UV-light for 2 minutes (Figure 3a). hMSC chondrogenic and osteogenic differentiation was accomplished by simultaneously introducing CM and OM into each separate chamber of the bioreactor (Figure 3b).

**Figure 3.**
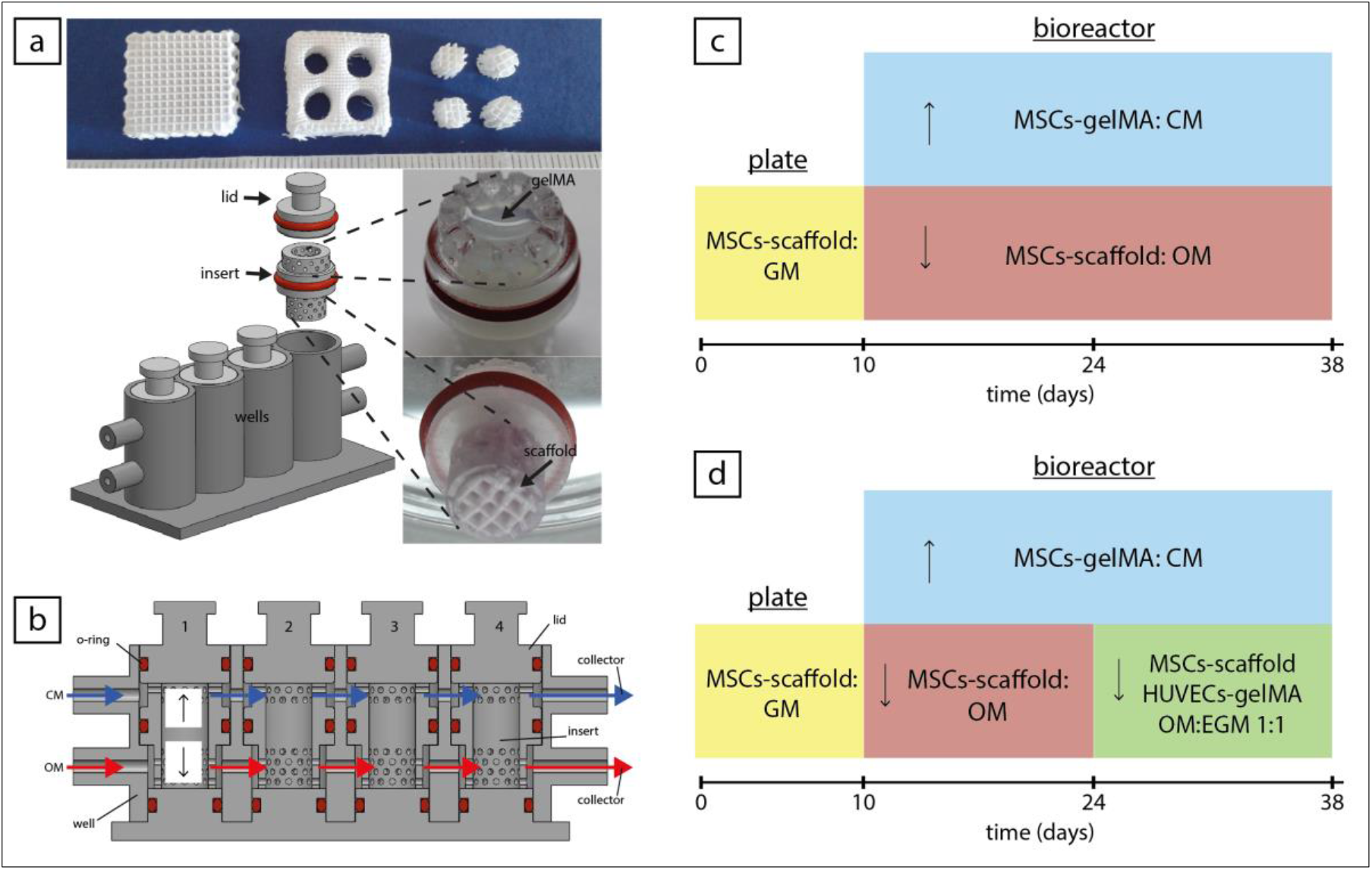
Osteochondral constructs preparation within the bioreactor and experimental design. (a) Cutting of a PCL scaffold to obtain four cylindrical samples and 3D visualization of the bioreactor with a wet-spun scaffold placed in the bottom part of the insert and gelMA placed in the upper part of the insert. (b) Schematic cross-section of the bioreactor used for the development of OC constructs, with chondrogenic and osteogenic media flow through the upper and lower part of the inserts, respectively. This configuration allowed the serial analysis of four constructs, the O-rings prevented medium leaking to the outside of the bioreactor and between upper and lower chambers, and the lids sealed each well to avoid contamination. (c) and (d) Experimental timelines for the development of biphasic OC constructs and triphasic vascularized OC constructs, respectively. GM = growth medium, OM = osteogenic medium, CM = chondrogenic medium, EGM = endothelial growth medium.

Figure 3c,d illustrate the experimental timelines and media composition for the generation of biphasic (chondro-osteo) and triphasic (chondro-osteo-vascular) constructs. After 4 weeks of differentiation, the biphasic constructs were removed from the insert (Figure 4, macroscopic) and analyzed to assess the chondrogenic and osteogenic differentiation of hMSCs. Macroscopic appearance of the engineered constructs resembled that of an OC plug removed from an articular joint, with the two phases securely bound to each other and stable during handling. This is relevant in developing OC scaffolds; for example, previous strategies used to integrate the osseous and chondral phases had employed melt-based processes [42,43] or a PLGA/β-TCP compact layer between bone and cartilage as both connector and insulator [44]. In this study, the cells of the chondral and osseous halves of the construct are ultimately encapsulated in a continuous gel. Specifically, the partial penetration of the uncrosslinked gelMA for the chondral phase into the initial layers of the wet-spun scaffold of the osseous phase, when photocrosslinked, created a continuous, crosslinked transition within the construct.

**Figure 4.**
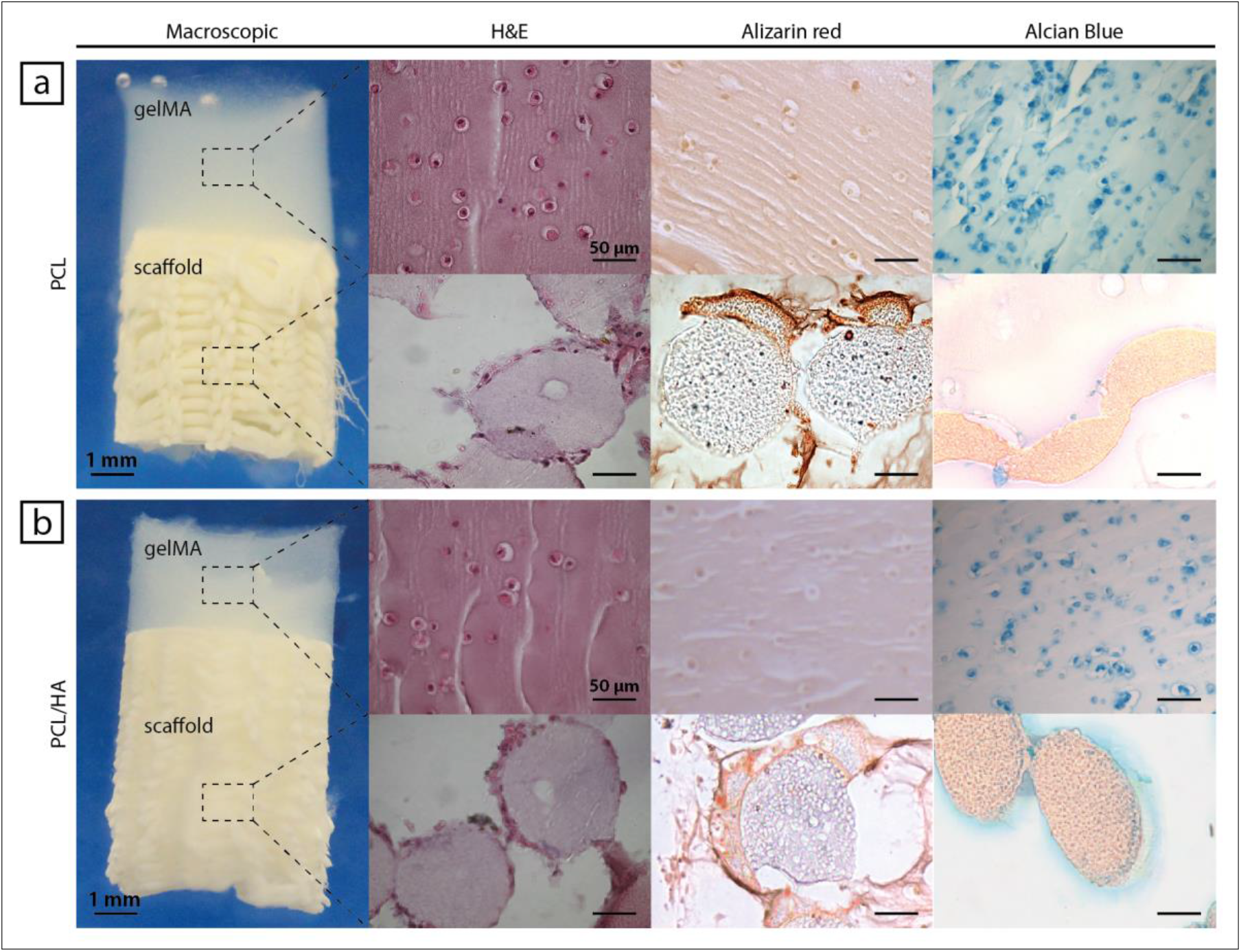
Non-vascularized biphasic osteochondral constructs with and without nanohydroxyapatite. Representative macroscopic images and histological analysis of the developed osteochondral constructs made of gelMA and PCL_1mm_ (a) or PCL/HA_1mm_ (b) scaffold seeded with hMSCs after 4 weeks of differentiation. Chondral component did not show calcium deposits by Alizarin red stain, while cells showed round-shape morphology by H&E stain (nuclei blue, cytoplasm pink-red). Osseous component did not show GAG depositions by Alcian blue stain, while cells appeared with a fusiform morphology by H&E stain.

Hematoxylin/eosin (H&E) staining showed the round-shaped morphology of hMSCs in the chondral construct, similar to the morphology of articular cartilage chondrocytes, while hMSCs in the osseous constructs presented the elongated shape typical of osteoblasts [45]. The biphasic constructs were sturdy during handling, showing mechanically strong cohesion between the cartilage phase and osseous phase. Histological analysis with Alcian blue/Alizarin red staining (Figure 4) revealed glycosaminoglycans (GAG) matrix in the upper chondral phase, concentrated inside lacunae-like structures around the cells, and calcium depositions in the osseous half, indicating distinct hMSCs differentiation into chondrogenic and osteogenic lineages. Notably, the gelMA-based phase did not show any calcium deposition, while the PCL-PCL/HA phase did not stain for Alcian blue (Figure 4), validating functional separation of the two differentiation media within the bioreactor.

### 3.3 Combination of engineered cartilage units and vascularized bone constructs in a triphasic model improved chondrogenesis and osteogenesis of MSCs

After evaluating the biphasic cartilage-bone only model, vascularized osseous constructs were combined with engineered cartilage to generate the triphasic model. As shown in Figure 5, histological analysis of the triphasic constructs showed separate staining of alcian blue and alizarin red in the gelMA and bone phase, respectively, highlighting distinct and adequate chondrogenic/osteogenic differentiation of the hMSCs. H&E staining confirmed the morphology of the differentiated hMSCs in chondrocytes and osteoblasts as shown in the biphasic constructs. The main innovation of our system is the introduction of endothelial cells together with hMSCs committed to chondrogenic and osteogenic differentiation. The presence of HUVEC-derived capillary-like networks was documented by the presence of positive staining for CD31 in the osseous compartments, while no CD31 positive stain were detected in the chondral phase.

**Figure 5.**
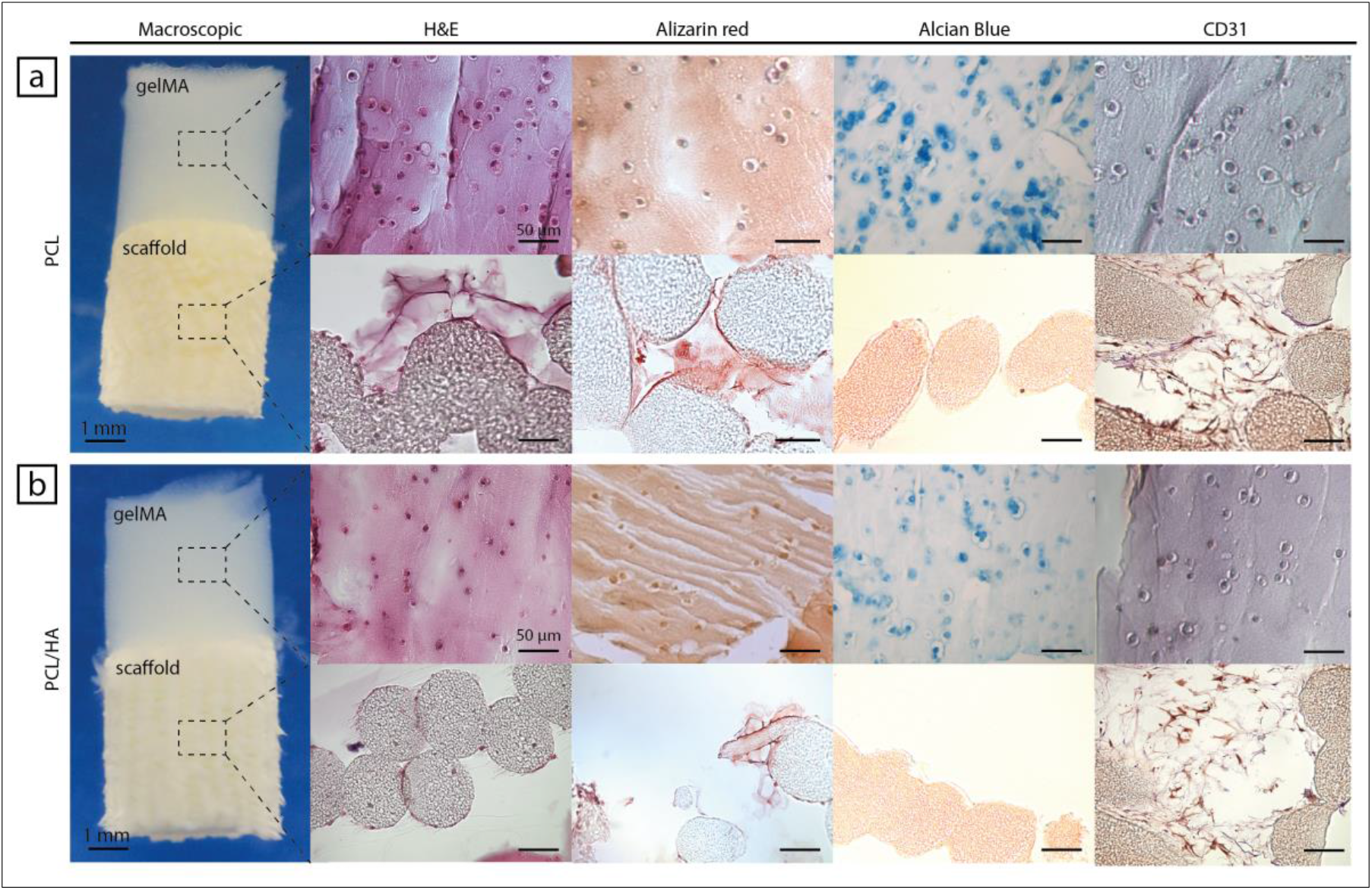
Tri-phasic vascularized osteochondral constructs with and without nanohydroxyapatite. Representative macroscopic images and histological/immunohistochemycal analysis of the developed vascularized osteochondral constructs made of gelMA and PCL_1mm_ (a) or PCL/HA_1mm_ (b) scaffold seeded with hMSCs after 4 weeks of differentiation. Chondral component did not show calcium deposits by Alizarin red stain, while cells exhibited round-shape morphology by H&E stain (nuclei blue, cytoplasm pink-red). Osseous component did not show GAG depositions by Alcian blue stain, while cells showed a fusiform morphology by H&E stain. CD31 immunohistochemycal analysis showed the presence of HUVECs in the PCL-PCL/HA phase, while no CD31 staining was seen in the corresponding gelMA parts.

The enhancement of chondrogenic/osteogenic differentiation of hMSCs by endothelial cells was quantified by gene expression analysis in the chondral and osseous halves via qRT-PCR. Cells seeded in the osseous part (PCL-PCL/HA scaffold) of the biphasic and triphasic constructs expressed higher level of osteogenic genes (OSC, OPN and BSP2) compared to day 0 undifferentiated cells, except for BSP2 in PCL/HA constructs (Figure 6), and osteogenic marker expression was increased more than 2-fold in the triphasic constructs as compared to biphasic ones. This finding demonstrates that the presence of endothelial cells enhanced osteogenic differentiation of hMSCs in the bioreactor system. Similarly, cells seeded in the chondral component (gelMA) upregulated chondral genes (COL2, SOX9, and ACAN) and downregulated osteogenic genes (OSC, OPN) compared to day 0 undifferentiated cells (Figure 7). The addition of HUVECs in the triphasic constructs significantly increased chondrogenic gene expression in the PCL constructs, but not in the PCL/HA constructs, a result supported by stronger Alcian blue staining in the triphasic constructs (Figure 5) compared to biphasic ones (Figure 4).

**Figure 6.**
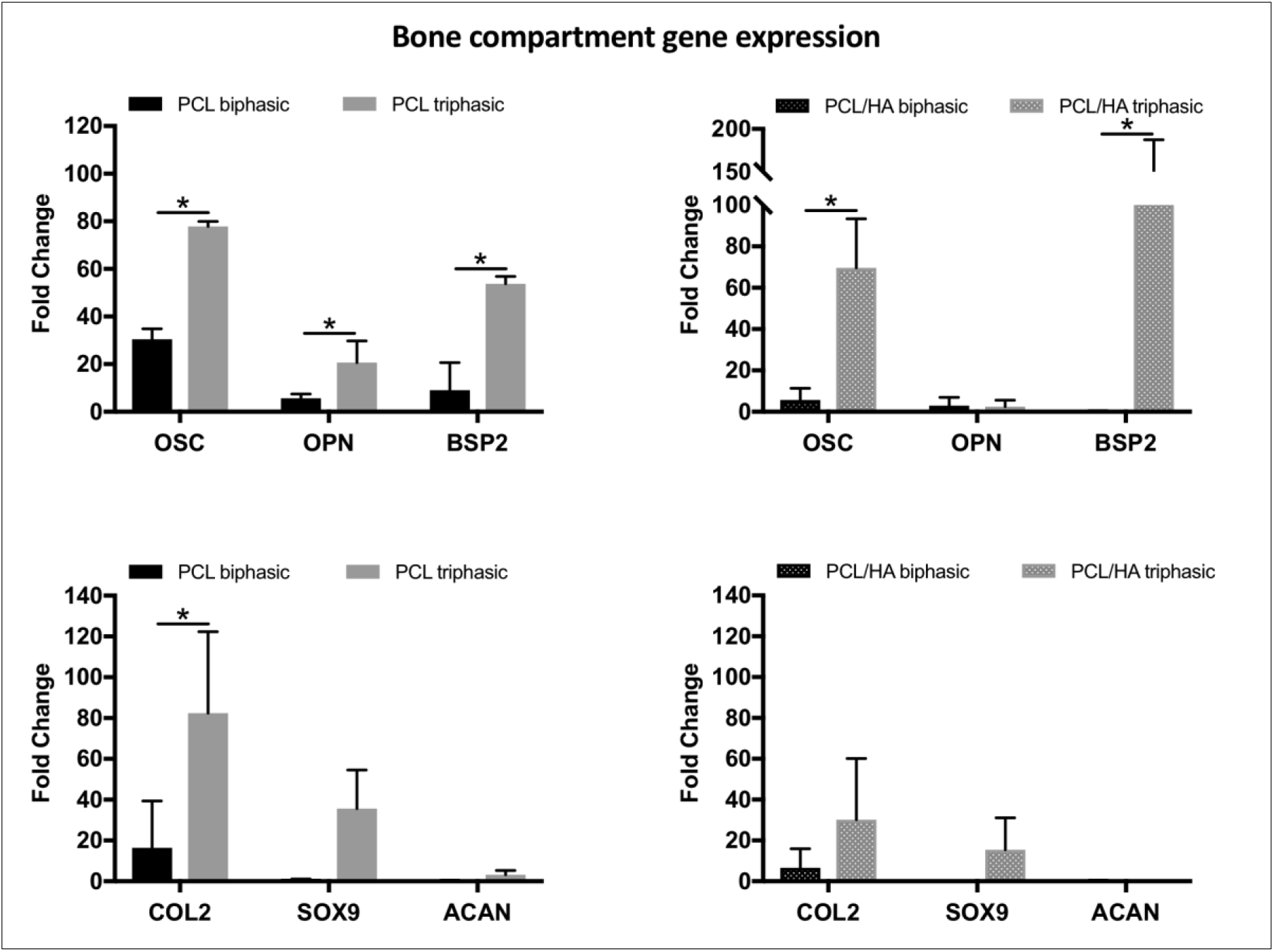
Analysis of gene expression in the osseous component for the assessment of osteogenic (OSC, OPN, BSP2) and chondrogenic (COL2, SOX9, ACAN) differentiation of hMSCs in PCL and PCL/HA scaffolds in the biphasic and triphasic constructs. Fold change = 1 represents the baseline of undifferentiated cells. Experiments were conducted on cells extracted from nine different patients divided in three groups of three (n = 3).

**Figure 7:**
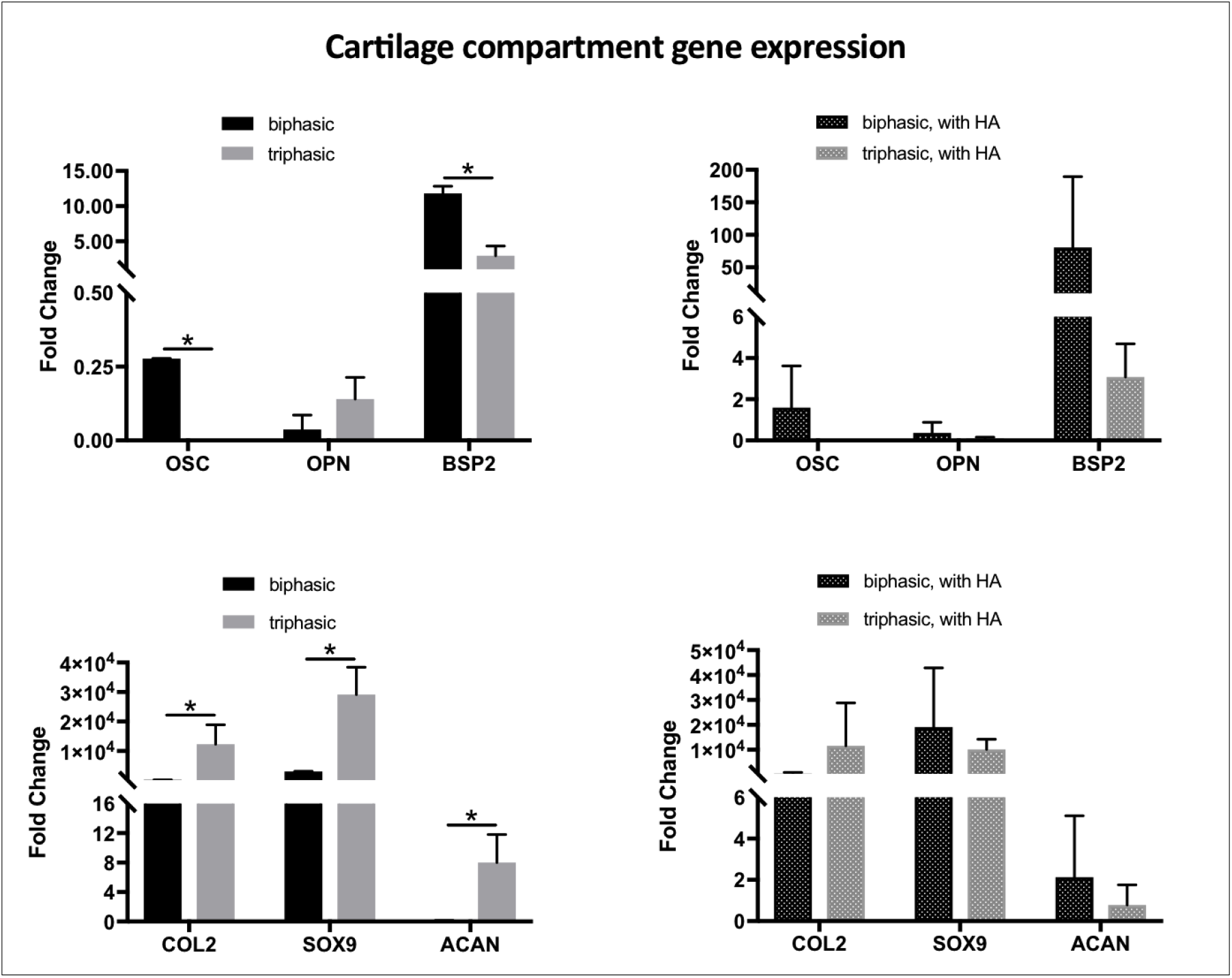
Analysis of gene expression in the cartilaginous component for osteogenic (OSC, OPN, BSP2) and chondrogenic (COL2, SOX9, ACAN) differentiation of hMSCs in gelMA in the biphasic and triphasic constructs. Fold change = 1 represents the baseline of undifferentiated cells. Experiments were conducted on cells extracted from nine different patients divided in three groups of three (n = 3).

This data also shows that the addition of HUVECs to the osseous component had no negative influence upon the adjacent engineered cartilage. We did observe a statistically significant but minimal increase in COL2 and SOX9 expression in the osseous component that is possibly related to low-level seepage of the chondrogenic medium from the top part of the construct, facilitated in part by the larger pores of the scaffold. Interestingly, the presence of the chondral component in the biphasic and triphasic constructs also had a positive effect on the expression of osteogenic genes compared to hMSCs alone and to hMSCs-HUVECs co-culture (compare Figures 2 and 6). Taken together, these data confirm the adequate functional separation of the two tissue-specific media and the ability of hMSCs to differentiate down these the two lineages.

One possible mechanism by which HUVECs could enhance chondrogenesis and osteogenesis could be related to the endothelin-1 (ET-1) protein secreted by endothelial cells as previously suggested by Tsai *et al.* [46]. We therefore examined ET-1 expression in our constructs. The results showed upregulation (Figure 8a) in the triphasic PCL-based constructs compared to triphasic PCL/HA constructs and to vascularized osseous constructs alone (PCL and PCL/HA scaffolds cultured in plate as described in Figure 2). As ET-1 has been postulated to act via activation of the AKT pathway [46], we also analyzed the presence of phosphorylated AKT (pAKT) through IHC. The results showed the presence of positive signals of pAKT in the triphasic constructs only, with stronger signals in the bone compartment and slightly less in the presence of HA (Figure 8b,c), corroborating the reduced expression of ET-1 in the presence of HA (Figure 8a).

**Figure 8.**
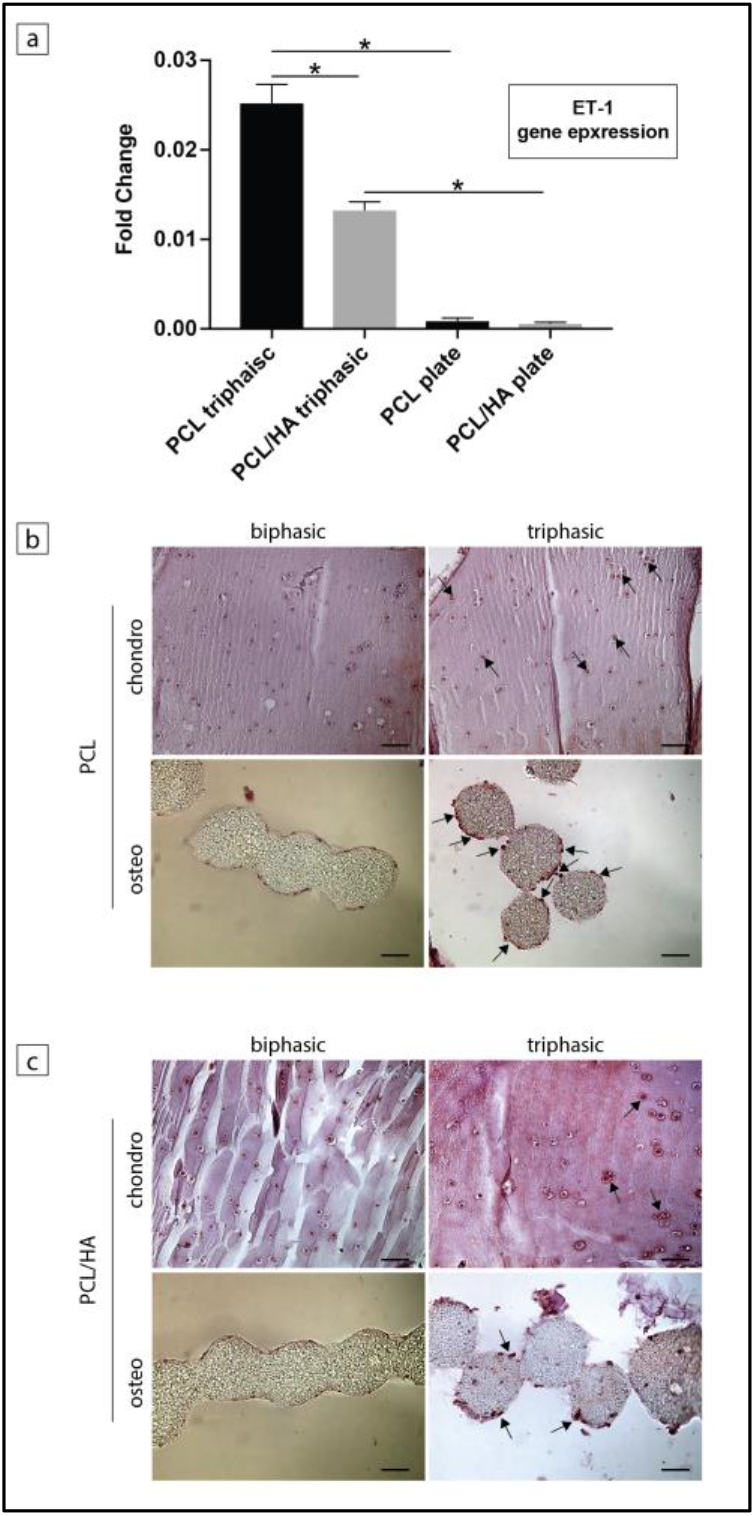
(a) Endothelin-1 (ET-1) gene expression in MPS bioreactor constructs vs. constructs engineered in plate. (b) and (c) representative IHC pictures of PCL and PCL/HA based constructs respectively, stained with anti-pAKT antibody: positive red signals are pointed by arrows. Scale bar = 100 μm

## 4 Discussion

The objective of this study was the development of an *in vitro* OC tissue suitable for the analysis of the cross-talk among the different cellular components involved in the homeostasis and patho-biology of joint diseases. A key issue in developing engineered osteochondral tissue is the introduction of a stable vasculature inside the bone construct [47]. This is hindered by technical and biological obstacles due to the differences between osteoblasts and endothelial cells. Different co-culture strategies have been exploited for the development of vascularized bone but, to the best of authors’ knowledge, the generation of a biomimetic vascularized OC unit has not been reported. The material choice for bone scaffolds gravitated toward PCL for its easy processability, biocompatibility and long history of successful application in bone tissue engineering applications. We also loaded the scaffolds with HA, since this is a mineral naturally present in bones and has been shown to support osteoconductivity *in vitro*. CAWS fabrication technique was chosen to produce bone scaffolds with structural integrity and interconnected porosity to allow growth and differentiation of hMSCs, while providing an entry point, a 3D space and a physical backbone for the introduction of HUVECs suspended in the gelMA hydrogel. The generation of such integrated construct can also enhance the functionalities of the resulting product, by combining the load-bearing properties and osteoconductivity of the developed wet-spun scaffold and the absorbing and mass transfer properties of the hydrogel [33]. GelMA has been previously demonstrated to be suitable for cartilage engineering, providing a 3D environment for hMSCs and ECM clues to allow a proper differentiation toward a chondrogenic phenotype [48].

Bone constructs differentiated in a plate showed, by both histology/immunohistochemistry and gene expression, enhanced osteogenic differentiation when the hMSCs seeded in the wet-spun scaffolds were co-cultured with HUVECs, which formed a capillary-like network. These results constitute the rationale for the integration of the vascularized engineered bone constructs within an OC biphasic model, with the purpose of developing a triphasic *in vitro* model of the OC complex that could better mimic native tissue physiology and serve as a platform for pathology studies and drug screening. The presence of hydroxyapatite (HA) reduced the enhancement of osteogenic differentiation in plate, with no positive effect in the case of OPN and OSC; furthermore, we observed slightly lower calcium deposits and less branched HUVEC-derived network. This phenomenon is in line with our previous results on the *in vitro* osteoconductivity of branched three-arm star PCL (*PCL)/HA scaffolds produced by CAWS and seeded with pre-osteoblasts [49], as well as our *in vivo* study on *PCL/HA scaffolds used for the regeneration of a critical size bone defect in a rabbit model [50]. HA nanoparticles can be internalized by HUVECs and accumulate in the cytoplasm, thus inducing dysfunction in the endothelial cells; this effect could explain the reduction in the enhancement of osteogenesis [51]. No significant changes in the expression of VEGF have been documented, whereas HA was revealed to interfere with angiogenesis by inhibiting the phosphorylation of endothelial nitric oxide synthase (eNOS) with a parallel downregulation of PI3K/AKT pathway [52]. These effects have been observed in our study, as seen by slightly less-branched HUVEC tubes (Figures 5b) and lower signals of phosphorylated AKT (Figure 8c) in the PCL/HA triphasic constructs.

The generation of a human cell-based, multi-tissue analog was made possible by the use of a next-generation bioreactor, which can help to recapitulate the complexity of biochemical interactions of human cells/tissues within the same fluidic system. The MPS OC bioreactor used in this work fits perfectly with the need of addressing simultaneous differentiation of different cell types toward specific tissues, while allowing the control of the local tissue specific microenvironments, which were optimized for hMSCs and HUVECs proliferation and differentiation. The bioreactor fluidics is designed to favor the separation of the upper and lower fluid flow and to minimize mixing [40]. This system was able to maintain two separate flows while, at the same time, allowing communication between the two phases that are in direct contact (Figure 4-5). By exploiting the excellent fluidics of the MPS bioreactor, we combined cartilage, bone and endothelial components in a unique construct. Of interest, osteogenesis of hMSCs was improved just by putting the scaffolds in the bioreactor with the presence of the chondrogenic gelMA component, as the level of osteogenic gene expression was almost doubled in the biphasic construct than bone scaffolds in plate (Figure 2b and Figure 6). This enhancement was even better in the triphasic model, with the HUVEC-derived capillary-like network present throughout 14 days of culture inside the bioreactor, showing a synergistic effect by the simultaneous presence of all cell types (Figure 6). Tsai *et al.* have previously demonstrated the enhancement of chondro- and osteogenesis of hMSCs in the presence of human aortic endothelial cells in 2D cultures [46]. In our model, as shown by histology and gene expression analysis (Figure 5-7), we documented the simultaneous enhancement of hMSC osteogenesis and chondrogenesis in a 3D multi-cellular environment due to the presence of HUVECs, which may or may not be in direct contact with hMSCs. A strong physical interconnection between the engineered tissue components was evident, supporting the feasibility of communication between the chondral and osseous components of the OC complex, as it occurs *in vivo*, and demonstrating the ability of our constructs to mimic a physiological environment.

We preliminarily investigated one of the possible mechanisms involved in this cross-talk among hMSCs and endothelial cells. AKT pathway has already been shown to be involved in the increased hMSC chondrogenesis/osteogenesis seen in the presence of HUVECs [46,53,54], as well as in several aspects of cell behavior, including osteoblast differentiation and bone development [55–57]. The presence of HUVECs resulted in the phosphorylation of AKT receptor both in the bone compartment and, at a lower extent, in the cartilage compartment, suggesting the communication between the two components (Figure 8). In particular, hMSCs in the bone compartment were always in close contact with HUVECs and specifically at the end of the 4-week culture period, while hMSCs in the cartilage compartment could experience a delayed effect, not significant at the 4-week culture end-point. In the future, a time-dependent activation of the pathway needs to be investigated to address better the effect of the HUVECs on hMSCs of both compartments through ET-1 mediated activation of AKT pathway.

Taken together, the combination of an engineered vascularized bone and accompanying cartilage as a single construct in a dual-flow bioreactor resulted in obvious synergistic effect in enhancing both chondrogenesis and osteogenesis of hMSCs, compared to constructs cultured separately, i.e., vascularized bone without cartilage and OC constructs without endothelial cells. The system developed here allowed analysis of the cross-talk among different cellular components and better elucidation of the interactions among the three cell types combined in a unique construct. Since these interactions are intimately involved in the development of the OC interface and the pathogenesis of degenerative joint diseases, our triphasic model represents a potential highly useful platform to study these phenomena.

## 5 Conclusions

The development of veritable *in vitro* models of the osteochondral unit is essential in understanding the biology of cartilage and bone development/regeneration, as well as for high throughput screening approaches for drug development. In this study, we successfully produced a triphasic construct by the combination of additively manufactured wet-spun PCL-PCL/HA scaffolds with gelMA. The engineering of these matrices encapsulating hMSCs and HUVECs, and the use a custom-designed MPS bioreactor, allowed the production of a vascularized OC construct characterized by satisfactory differentiation towards both chondral and osseous phenotypes, in term of GAG/calcium deposition and gene expression. Endothelial cells showed the ability to form capillary-like networks and enhanced the osteogenic and chondrogenic differentiation of the hMSCs in both compartments, as compared to constructs without HUVECs. The synergistic enhancement of chondrogenesis and osteogenesis in our triphasic model represents a promising step towards the development of a veritable model of the OC complex, applicable for the study of osteochondral pathologies and drug screening.

## Acknowledgments

The authors gratefully acknowledge Luca Coluccino, Ph.D. for providing the gelMA, and Jian Tan, M.D. for isolation of hMSCs. This work was supported in part by the *Ministero dell’Istruzione*, *dell’Università e della Ricerca* (DM186/2008), Ri.MED Foundation (Italy), the U.S. Department of Defense (W81XWH-08-2-0032 and W81XWH-10-1-0850), and the National Institutes of Health (NCATS 1UG3 TR002136 and NIAMS 1R43AR072169-01).

## References

[1] M. Lethbridge-Cejku, C.G. Helmick, J.R. Popovic, Hospitalizations for arthritis and other rheumatic conditions: data from the 1997 National Hospital Discharge Survey., Med. Care. 41 (2003) 1367–73.

[2] A. Carbone, S. Rodeo, Review of current understanding of post-traumatic osteoarthritis resulting from sports injuries, J. Orthop. Res. 35 (2017) 397–405.

[3] K.D. Illingworth, D. Hensler, B. Casagranda, C. Borrero, C.F. van Eck, F.H. Fu, Relationship between bone bruise volume and the presence of meniscal tears in acute anterior cruciate ligament rupture, Knee Surg. Sports Traumatol. Arthrosc. 22 (2014) 2181–2186.

[4] L. Zhang, J.D. Hacke, W.E. Garrett, H. Liu, B. Yu, Bone Bruises Associated with Anterior Cruciate Ligament Injury as Indicators of Injury Mechanism: A Systematic Review, Sport. Med. 49 (2019) 453–462.

[5] C. Kia, Z. Cavanaugh, E. Gillis, C. Dwyer, V. Chadayammuri, L.N. Muench, D.P. Berthold, M. Murphy, R. Pacheco, R.A. Arciero, Size of Initial Bone Bruise Predicts Future Lateral Chondral Degeneration in ACL Injuries: A Radiographic Analysis, Orthop. J. Sport. Med. 8 (2020) 232596712091683.

[6] M. Calvo-Gurry, E.T. Hurley, D. Withers, M. Vioreanu, R. Moran, Posterior tibial bone bruising associated with posterior-medial meniscal tear in patients with acute anterior cruciate ligament injury, Knee Surgery, Sport. Traumatol. Arthrosc. 27 (2019) 3633–3637.

[7] O.H. Sandberg, P. Aspenberg, Inter-trabecular bone formation: a specific mechanism for healing of cancellous bone: A narrative review, Acta Orthop. 87 (2016) 459–465.

[8] R.J. Lories, F.P. Luyten, The bone-cartilage unit in osteoarthritis., Nat. Rev. Rheumatol. 7 (2011) 43–9.

[9] S. Lopa, H. Madry, Bioinspired scaffolds for osteochondral regeneration., Tissue Eng. Part A. 20 (2014) 2052–76.

[10] P. Nooeaid, V. Salih, J.P. Beier, A.R. Boccaccini, Osteochondral tissue engineering: scaffolds, stem cells and applications., J. Cell. Mol. Med. 16 (2012) 2247–70.

[11] A. Di Luca, C. Van Blitterswijk, L. Moroni, The osteochondral interface as a gradient tissue: From development to the fabrication of gradient scaffolds for regenerative medicine, Birth Defects Res. Part C Embryo Today Rev. 105 (2015) 34–52.

[12] C. Sanchez, M.A. Deberg, N. Piccardi, P. Msika, J.-Y.L. Reginster, Y.E. Henrotin, Subchondral bone osteoblasts induce phenotypic changes in human osteoarthritic chondrocytes, Osteoarthr. Cartil. 13 (2005) 988–997.

[13] M.B. Goldring, S.R. Goldring, Articular cartilage and subchondral bone in the pathogenesis of osteoarthritis, Ann. N. Y. Acad. Sci. 1192 (2010) 230–237.

[14] S. Ashraf, P.I. Mapp, D.A. Walsh, Contributions of angiogenesis to inflammation, joint damage, and pain in a rat model of osteoarthritis, Arthritis Rheum. 63 (2011) 2700–2710.

[15] M. Saito, T. Sasho, S. Yamaguchi, N. Ikegawa, R. Akagi, Y. Muramatsu, S. Mukoyama, N. Ochiai, J. Nakamura, K. Nakagawa, A. Nakajima, K. Takahashi, Angiogenic activity of subchondral bone during the progression of osteoarthritis in a rabbit anterior cruciate ligament transection model, Osteoarthr. Cartil. 20 (2012) 1574–1582.

[16] S. Suri, S.E. Gill, S. Massena de Camin, D. Wilson, D.F. McWilliams, D.A. Walsh, Neurovascular invasion at the osteochondral junction and in osteophytes in osteoarthritis., Ann. Rheum. Dis. 66 (2007) 1423–8.

[17] J. Harper, M. Klagsbrun, Cartilage to bone - Angiogenesis leads the way, Nat. Med. 5 (1999) 617–618.

[18] M.-H. Lafage-Proust, B. Roche, M. Langer, D. Cleret, A. Vanden Bossche, T. Olivier, L. Vico, Assessment of bone vascularization and its role in bone remodeling, Bonekey Rep. 4 (2015) 662.

[19] K.D. Hankenson, M. Dishowitz, C. Gray, M. Schenker, Angiogenesis in bone regeneration, Injury. 42 (2011) 556–561.

[20] S.A. Wong, K.O. Rivera, T. Miclau, E. Alsberg, R.S. Marcucio, C.S. Bahney, Microenvironmental Regulation of Chondrocyte Plasticity in Endochondral Repair—A New Frontier for Developmental Engineering, Front. Bioeng. Biotechnol. 6 (2018) 58.

[21] Y. Kawakami, T. Matsumoto, Y. Mifune, T. Fukui, K. Patel, G. Walker, M. Kurosaka, R. Kuroda, Therapeutic Potential of Endothelial Progenitor Cells in the Field of Orthopaedics, Curr. Stem Cell Res. Ther. 12 (2016) 3–13.

[22] A. Pirosa, R. Gottardi, P.G. Alexander, R.S. Tuan, Engineering in-vitro stem cell-based vascularized bone models for drug screening and predictive toxicology, Stem Cell Res. Ther. 9 (2018).

[23] Y.H. Shen, M.S. Shoichet, M. Radisic, Vascular endothelial growth factor immobilized in collagen scaffold promotes penetration and proliferation of endothelial cells., Acta Biomater. 4 (2008) 477–89.

[24] E.A. Bayer, J. Jordan, A. Roy, R. Gottardi, M.V. Fedorchak, P.N. Kumta, S.R. Little, Programmed platelet-derived growth factor-bb and bone morphogenetic protein-2 delivery from a hybrid calcium phosphate/alginate scaffold, Tissue Eng. -Part A. 23 (2017).

[25] Y. Liu, S.-H. Teoh, M.S.K. Chong, E.S.M. Lee, C.N.Z. Mattar, N.K. Randhawa, Z.-Y. Zhang, R.J. Medina, R.D. Kamm, N.M. Fisk, M. Choolani, J.K.Y. Chan, Vasculogenic and osteogenesis-enhancing potential of human umbilical cord blood endothelial colony-forming cells., Stem Cells. 30 (2012) 1911–24.

[26] O. Tsigkou, I. Pomerantseva, J.A. Spencer, P.A. Redondo, A.R. Hart, E. O’Doherty, Y. Lin, C.C. Friedrich, L. Daheron, C.P. Lin, C.A. Sundback, J.P. Vacanti, C. Neville, Engineered vascularized bone grafts, Proc. Natl. Acad. Sci. 107 (2010) 3311–3316.

[27] I. Chiesa, C. De Maria, A. Lapomarda, G.M. Fortunato, F. Montemurro, R. Di Gesù, R.S. Tuan, G. Vozzi, R. Gottardi, Endothelial cells support osteogenesis in an in vitro vascularized bone model developed by 3D bioprinting, Biofabrication. 12 (2020) 25013.

[28] P.G. Alexander, R. Gottardi, H. Lin, T.P. Lozito, R.S. Tuan, Three-dimensional osteogenic and chondrogenic systems to model osteochondral physiology and degenerative joint diseases, Exp. Biol. Med. 239 (2017) 1080–1095.

[29] T.P. Lozito, P.G. Alexander, H. Lin, R. Gottardi, A.W.-M. Cheng, R.S. Tuan, Three-dimensional osteochondral microtissue to model pathogenesis of osteoarthritis., Stem Cell Res. Ther. 4 Suppl 1 (2013) S6.

[30] J.A. Benton, C.A. DeForest, V. Vivekanandan, K.S. Anseth, Photocrosslinking of Gelatin Macromers to Synthesize Porous Hydrogels That Promote Valvular Interstitial Cell Function, Tissue Eng. Part A. 15 (2009) 3221–3230.

[31] D. Puppi, C. Mota, M. Gazzarri, D. Dinucci, A. Gloria, M. Myrzabekova, L. Ambrosio, F. Chiellini, Additive manufacturing of wet-spun polymeric scaffolds for bone tissue engineering., Biomed. Microdevices. 14 (2012) 1115–27.

[32] D.A. Nichols, I.S. Sondh, S.R. Litte, P. Zunino, R. Gottardi, Design and validation of an osteochondral bioreactor for the screening of treatments for osteoarthritis, Biomed. Microdevices. 20 (2018).

[33] D. Puppi, C. Migone, L. Grassi, A. Pirosa, G. Maisetta, G. Batoni, F. Chiellini, Integrated 3D Fibers/Hydrogel Biphasic Scaffolds for Periodontal Bone Tissue Engineering, Polym. Int. (2016) n/a-n/a.

[34] D. Puppi, A.M. Piras, A. Pirosa, S. Sandreschi, F. Chiellini, Levofloxacin-loaded star poly(ε-caprolactone) scaffolds by additive manufacturing., J. Mater. Sci. Mater. Med. 27 (2016) 44.

[35] A.I. Van Den Bulcke, B. Bogdanov, N. De Rooze, E.H. Schacht, M. Cornelissen, H. Berghmans, Structural and rheological properties of methacrylamide modified gelatin hydrogels., Biomacromolecules. 1 (2000) 31–8.

[36] B.D. Fairbanks, M.P. Schwartz, C.N. Bowman, K.S. Anseth, Photoinitiated polymerization of PEG-diacrylate with lithium phenyl-2,4,6-trimethylbenzoylphosphinate: polymerization rate and cytocompatibility., Biomaterials. 30 (2009) 6702–7.

[37] E.J. Caterson, L.J. Nesti, K.G. Danielson, R.S. Tuan, Human Marrow-Derived Mesenchymal Progenitor Cells: isolation, culture expansion, and analysis of differentiation, Mol. Biotechnol. 20 (2002) 245–256.

[38] T.O. Pedersen, A.L. Blois, Z. Xing, Y. Xue, Y. Sun, A. Finne-Wistrand, L.A. Akslen, J.B. Lorens, K.N. Leknes, I. Fristad, K. Mustafa, Endothelial microvascular networks affect gene-expression profiles and osteogenic potential of tissue-engineered constructs., Stem Cell Res. Ther. 4 (2013) 52.

[39] R.R. Rao, M.L. Vigen, A.W. Peterson, D.J. Caldwell, A.J. Putnam, J.P. Stegemann, Dual-Phase Osteogenic and Vasculogenic Engineered Tissue for Bone Formation, Tissue Eng. Part A. 21 (2014) 530–540.

[40] L. Iannetti, G. D’Urso, G. Conoscenti, E. Cutrì, R.S. Tuan, Distributed and Lumped Parameter Models for the Characterization of High Throughput Bioreactors, PLoS One. 11 (2016) e0162774.

[41] H. Lin, T.P. Lozito, P.G. Alexander, R. Gottardi, R.S. Tuan, Stem cell-based microphysiological osteochondral system to model tissue response to interleukin-1β., Mol. Pharm. 11 (2014) 2203–12.

[42] S. Ghosh, J.C. Viana, R.L. Reis, J.F. Mano, Bi-layered constructs based on poly(l-lactic acid) and starch for tissue engineering of osteochondral defects, Mater. Sci. Eng. C. 28 (2008) 80–86.

[43] J.E. Jeon, C. Vaquette, C. Theodoropoulos, T.J. Klein, D.W. Hutmacher, Multiphasic construct studied in an ectopic osteochondral defect model., J. R. Soc. Interface. 11 (2014) 20140184.

[44] H. Da, S.-J. Jia, G.-L. Meng, J.-H. Cheng, W. Zhou, Z. Xiong, Y.-J. Mu, J. Liu, The impact of compact layer in biphasic scaffold on osteochondral tissue engineering., PLoS One. 8 (2013) e54838.

[45] E. Beltrán-Partida, A. Moreno-Ulloa, B. Valdez-Salas, C. Velasquillo, M. Carrillo, A. Escamilla, E. Valdez, F. Villarreal, Improved Osteoblast and Chondrocyte Adhesion and Viability by Surface-Modified Ti6Al4V Alloy with Anodized TiO2 Nanotubes Using a Super-Oxidative Solution, Materials (Basel). 8 (2015) 867–883.

[46] T.-L. Tsai, B. Wang, M.W. Squire, L.-W. Guo, W.-J. Li, Endothelial cells direct human mesenchymal stem cells for osteo- and chondro-lineage differentiation through endothelin-1 and AKT signaling., Stem Cell Res. Ther. 6 (2015) 88.

[47] Y. Liu, J.K.Y. Chan, S.-H. Teoh, Review of vascularised bone tissue-engineering strategies with a focus on co-culture systems, J. Tissue Eng. Regen. Med. 9 (2015) 85–105.

[48] W. Schuurman, P.A. Levett, M.W. Pot, P.R. van Weeren, W.J.A. Dhert, D.W. Hutmacher, F.P.W. Melchels, T.J. Klein, J. Malda, Gelatin-Methacrylamide Hydrogels as Potential Biomaterials for Fabrication of Tissue-Engineered Cartilage Constructs, Macromol. Biosci. 13 (2013) 551–561.

[49] C. Mota, D. Puppi, D. Dinucci, M. Gazzarri, F. Chiellini, Additive manufacturing of star poly(- caprolactone) wet-spun scaffolds for bone tissue engineering applications, J. Bioact. Compat. Polym. 28 (2013) 320–340.

[50] F. Dini, G. Barsotti, D. Puppi, A. Coli, A. Briganti, E. Giannessi, V. Miragliotta, C. Mota, A. Pirosa, M.R. Stornelli, P. Gabellieri, F. Carlucci, F. Chiellini, Tailored star poly (- caprolactone) wet-spun scaffolds for in vivo regeneration of long bone critical size defects, J. Bioact. Compat. Polym. 31 (2016) 15–30.

[51] X. Liu, J. Sun, Potential proinflammatory effects of hydroxyapatite nanoparticles on endothelial cells in a monocyte-endothelial cell coculture model., Int. J. Nanomedicine. 9 (2014) 1261–73.

[52] X. Shi, K. Zhou, F. Huang, C. Wang, Interaction of hydroxyapatite nanoparticles with endothelial cells: internalization and inhibition of angiogenesis in vitro through the PI3K/Akt pathway, Int. J. Nanomedicine. Volume 12 (2017) 5781–5795.

[53] D. Ikegami, H. Akiyama, A. Suzuki, T. Nakamura, T. Nakano, H. Yoshikawa, N. Tsumaki, Sox9 sustains chondrocyte survival and hypertrophy in part through Pik3ca-Akt pathways, Development. 138 (2011) 1507–1519.

[54] L. Ling, C. Dombrowski, K.M. Foong, L.M. Haupt, G.S. Stein, V. Nurcombe, A.J. van Wijnen, S.M. Cool, Synergism between Wnt3a and Heparin Enhances Osteogenesis via a Phosphoinositide 3-Kinase/Akt/RUNX2 Pathway, J. Biol. Chem. 285 (2010) 26233–26244.

[55] I.M. McGonnell, A.E. Grigoriadis, E.W.-F. Lam, J.S. Price, A. Sunters, A specific role for phosphoinositide 3-kinase and AKT in osteoblasts?, Front. Endocrinol. (Lausanne). 3 (2012) 88.

[56] C.C. Mandal, G. Ghosh Choudhury, N. Ghosh-Choudhury, Phosphatidylinositol 3 Kinase/Akt Signal Relay Cooperates with Smad in Bone Morphogenetic Protein-2-Induced Colony Stimulating Factor-1 (CSF-1) Expression and Osteoclast Differentiation, Endocrinology. 150 (2009) 4989–4998.

[57] A. Mukherjee, P. Rotwein, Akt promotes BMP2-mediated osteoblast differentiation and bone development., J. Cell Sci. 122 (2009) 716–26.

